# Probing hippocampal stimulation in experimental temporal lobe epilepsy with functional MRI

**DOI:** 10.1101/2023.09.07.556678

**Authors:** Niels Schwaderlapp, Enya Paschen, Pierre LeVan, Dominik von Elverfeldt, Carola Haas

**Author notes:** Corresponding author: Dr. Niels Schwaderlapp, Medical Physics, Department of Radiology, University Medical Center Freiburg, Kilianstraße 5a, 79106 Freiburg, Germany.

## Abstract

**Background:** Electrical neurostimulation is a potentially effective therapy in epilepsy but the optimal approach is not yet clear. The parameter space is wide and the effects of different stimulations are not immediately obvious. Functional MRI (fMRI) can reveal which brain areas are affected by stimulation and help understand the induced effects. However, simultaneous deep brain stimulation (DBS)-fMRI examinations in patients are rare and the possibility to investigate multiple stimulation protocols is limited. Preclinical stimulation-fMRI studies can provide predictive value and help identify optimal neurostimulation parameters.

**Objective:** To systematically investigate the brain-wide responses to hippocampal electrical stimulations in a mouse model of mesial temporal lobe epilepsy (mTLE) using fMRI.

**Methods:** We applied electrical stimulation in the intrahippocampal kainate mouse model of mTLE and assessed the effect of different stimulation amplitudes (80-230 µA) and frequencies (1-100 Hz). In addition, the effect of prolonged 1 Hz stimulation was explored. Saline-injected mice served as controls.

**Results:** Varying the stimulation amplitudes had little effect on the resulting activation patterns. Low frequency stimulation led to a local response at the stimulation site only, whereas high frequency resulted in a spread of activation from the hippocampal formation into cortical and frontal areas. Prolonged low frequency stimulation reduced excitability.

**Conclusions:** While the amplitude parameter offers little opportunity to vary the outcome, the frequency represented the key parameter and determined whether the induced activation remained local or spread across the brain. This is in line with the few DBS-fMRI results obtained in epilepsy patients demonstrating the translational value of fMRI.

## 1. Introduction

Brain stimulation has become an established treatment option in movement disorders, such as Parkinson’s disease, and is increasingly becoming considered in the treatment of intractable epilepsy [1,2]. Temporal lobe epilepsy (TLE) in particular, the most common form of focal epilepsy, is often refractory to antiseizure medications. In these cases, the surgical resection of epileptic tissue is usually considered the best chance to attain seizure freedom, but there is no guarantee that surgery will result in a positive long-term outcome and will not be associated with post-operative cognitive deficits. Seizure recurrence is particularly high (20-50 %) when poor predictive criteria such as a long history of seizures or no identifiable focus are met [3–8]. This underscores the need to develop further treatment options.

The potential benefit of neurostimulation in epilepsy may be based on two mechanisms. First, a stimulation initiated at the onset of an epileptic seizure could desynchronize neuronal activity and thus terminate the seizure or at least prevent further propagation. Second, stimulation could lead to a reduction of neuronal excitability and thus reduce the probability of seizure occurrence. While there is promising evidence that stimulation has a positive therapeutic effect [9], it does not always achieve complete seizure freedom and is therefore rarely considered unless other avenues have been exhausted. A current challenge is to better investigate the effects of stimulation in order to increase effectiveness and minimize side effects.

The outcome is largely influenced by the stimulation site and the exact stimulation parameters. In mesial TLE (mTLE), one obvious target is the hippocampus (HC) [10–14]. In many patients, this is the presumed site of seizure onset and is therefore a frequent target for implanted electrodes during presurgical evaluation, enabling stimulation experiments to be performed. Initial stimulation parameters are selected based on experience and are adjusted to individual patients on a trial-and-error basis until a certain level of subjective behavioral response is achieved. If the response is sub-optimal or side effects occur, parameters such as amplitude or frequency are varied [15]. This time-consuming process is made more difficult by the fact that it is usually unknown which brain regions and networks are affected by the stimulation, which depends on the parameters as well as the individual disease condition. In addition, it is difficult to identify and adapt to long-term changes over time [16,17].

A valuable method for directly assessing the brain-wide responses to stimulation is functional MRI, which has been performed concurrently with deep brain stimulation (DBS) in recent studies [18–21]. However, due to the complexity of performing DBS and fMRI simultaneously, published reports from epilepsy patients remain scarce. For example, stimulation in the anterior nucleus of the thalamus showed a more widespread activation at 145 Hz than at 30 Hz stimulation [19]. In another study, similar activation patterns were observed when high (8 mA) and low (4 mA) currents were compared at 20 Hz stimulation, whereas no significant fMRI response was found in patients stimulated at 1 Hz [21]. Further studies are needed to better understand the influence of stimulation parameters. However, diverse patient cohorts and time constraints hamper systematic variation of parameters in clinical studies.

This investigation can be accelerated by preclinical DBS-fMRI studies. Studies targeting the rodent HC showed fMRI activations within the hippocampal structure and in additional areas, e.g. thalamus and septum, at stimulation frequencies of 10-130 Hz, but no fMRI response at 2 Hz electrical stimulation [22–24]. However, these studies were performed in healthy rodents. Therefore, it remains unclear to what extent these results, in particular the spread of activation from the HC to connected areas, reflect the situation in mTLE, which is associated with severe structural alterations of the HC [25–28].

The alterations of the HC that occur in the human disease are replicated particularly well in the intrahippocampal kainate (ihKA) model of mTLE [29–31]. After kainate injection and a latent phase, hippocampal sclerosis, characterized by segmental cell death and gliosis in the affected HC, is established and spontaneous seizures occur in the chronic stage. Results obtained from the ihKA model of mTLE can therefore be translated relatively well to human mTLE [32]. However, to our knowledge, hippocampal stimulation in this model has not been investigated with fMRI.

Non-imaging studies in experimental TLE examined the impact of electrical stimulation in the HC on the rate of epileptiform activity. Long-term (several days) high-frequency stimulation was found to be most effective in reducing seizure rates [33–35], but low frequencies (1–5 Hz) also showed promise [36–39]. The optimal approach to reduce seizure rates without causing severe side effects remains elusive.

In this study, we investigated the impact of multiple stimulation parameters in mTLE using fMRI. By stimulating the HC in the ihKA model of mTLE, we found that the frequency was the most important parameter and determined whether activity was confined to the hippocampal structure or spread across the brain. We further investigated the potential of prolonged 1 Hz stimulation to reduce the local excitability.

## 2. Materials and Methods

### Animals

Adult transgenic male mice (n=12, Cre-negative littermates from C57BL/6-Tg(Rbp4-cre)KL100Gsat x Ai32(RCL-ChR2(H134R)/EYFP) breeding) were used in this study. All animal procedures were carried out under the guidelines of the European Community’s Council Directive (2010/63/EU) and were approved by the regional council (Regierungspräsidium Freiburg, G-19/04).

### Kainate injections

Mice were injected with 50 nl kainate (Tocris, Bristol, UK) (n = 6) or saline (n = 6) into the right dorsal HC. In brief, the stereotaxic injection was performed under deep anesthesia (ketamine hydrochloride 100 mg/kg, xylazine 5 mg/kg, atropine 0.1 mg/kg body weight, i.p.) using Nanoject III (Drummond Scientific Company, Broomall, PA, USA). Mice were randomly assigned to be kainate-(15 mM in 0.9% sterile saline) or saline-injected. Stereotaxic coordinates relative to Bregma (in mm) were: anterioposterior (AP) = −2.0, mediolateral (ML) = −1.5, and relative to the cortical surface: dorsoventral (DV) = −1.5. Following kainate injection the occurrence of a behavioral status epilepticus was verified, characterized by mild convulsion, chewing, immobility, or rotations, as described before [40,41].

### Construction of electrodes

The electrodes were made of carbon to minimize susceptibility artifacts during MRI measurements [42,43]. For the stimulation/recording electrodes, sections of 1k carbon fiber tow (CST Composites, Tehachapi, California, USA), consisting of 1000 carbon filaments, were split five times to achieve bundles of approximately 0.2k. For the electrodes serving as ground/reference, four of these 1k tows were put together to achieve a lower current density and to prevent stimulation at the ground/reference electrodes. The carbon bundles were cold soldered to sections of copper wire using conductive silver epoxy (MG Chemicals, Ontario, Canada). The electrodes were coated with two layers of polydimethylsiloxane (PDMS, Sylgard 184, The Dow Chemical Company, USA) to ensure a biocompatible insulation. The carbon ends were cut to 3 mm length to expose the contacts prior to implantation.

### Electrode implantations

Sixteen days after kainate/saline injections, electrodes were implanted into the ipsilateral and contralateral dorsal HC (HCi and HCc, respectively). Stereotaxic coordinates are given relative to Bregma in mm for anterioposterior (AP) and mediolateral (ML) or to the cortical surface for dorsoventral (DV) coordinates: HCc: AP = −2.0, ML = +1.4, DV = −1.6 ; HCi: AP = −2.0, ML = −1.4, DV = −1.6.Two thick carbon fiber bundles were placed above the frontal cortex to provide reference and ground. The implants were fixed with dental cement (Paladur, Kulzer GmbH, Hanau, Germany).

### MRI acquisition

Anesthesia was initiated with ca. 5 % sevoflurane (AbbVie, Ludwigshafen, Germany) and maintained at ca. 2 % during setup. Subsequently, the anesthesia was switched to medetomidine (Domitor, Vetoquinol, Germany; s.c. bolus 0.3 mg/kg, followed by continuous s.c. infusion 0.6 mg/kg/h). Medetomidine, unlike isoflurane, does not suppress epileptiform activity [44,45]. During scans, animals were supplied with air (30 % O_2_), and respiration (ca.120-170 breaths/min) and body temperature (36 °C ± 1.5 °C) were monitored. After the scans, atipamezole (Antisedan, Vetoquinol, Germany) was administered (s.c., 0.8 mg/kg) to reverse the sedative effect of medetomidine.

MRI data were acquired on a preclinical 7 T system (BioSpec 70/20, Bruker BioSpin, Ettlingen, Germany) using a 2-channel surface transmit-receive cryoprobe. Adjustments included local shimming up to 3^rd^ order, adapted to the brain volume. An anatomical scan was acquired using a fast spin-echo sequence (RARE) with the parameters: TR 4 s, effective TE 40 ms, RARE factor 4, encoding matrix 120 x 64, in-plane resolution 0.14 x 0.14 mm^2^, number of slices 12, slice thickness 0.8 mm, slice gap 0.2 mm.

FMRI data were acquired using a gradient, multi-echo EPI sequence with the parameters: TR 1.5 s, TE_1_ 14 ms, TE_2_ 23 ms and TE_3_ 32 ms (three echoes), encoding matrix 60 x 32, in-plane resolution 0.28 x 0.28 mm^2^, number of slices 12, slice thickness 0.8 mm, slice gap 0.2 mm, excitation pulse flip-angle 70°, acquisition bandwidth 250 kHz.

### fMRI postprocessing

Data analysis was performed using FSL (FMRIB, Oxford, UK) and Matlab (Mathworks, USA). Preprocessing included motion correction, slice timing correction and highpass temporal filtering (0.01 Hz). The three individual time series (one for each echo) were combined into a single “multi-echo” dataset by T2*-weighted combination [46]. After manual brain extraction, the images were spatially smoothed using a Gaussian kernel with a full-width at half-maximum of 1.5 times in-plane resolution.

The activation was analyzed using the general linear model (GLM) in FSL [47,48]. In the first level, the stimulation timings were convolved with finite impulse response functions (8^th^ order, 24 s window) to create the design matrix [49]. In the group level GLM, a fixed-effects model was used and the results were registered to the AMBMC atlas space [50,51].

### Local field potential recordings

During fMRI measurements, local field potential (LFP) signals were recorded (EGI, Electrical Geodesics, USA) in the left dorsal HC with a sampling rate of 1 kHz. The data were analyzed with the toolbox EEGLAB [52] in Matlab (Mathworks, USA). Postprocessing included removal of the 50 Hz line noise, a moving average FIR filter (order 20) to remove residual line noise and MRI gradient artifacts, and temporal highpass filtering at 2 Hz.

### Electrical stimulations

The stimulation (STG1004, Smart Ephys/Multi-Channel Systems, Reutlingen, Germany) was performed through the electrode in the right dorsal HC with charged balanced bi-phasic pulses; 400µs phase duration, anodic first.

A single block-design experiment consisted of four 10 s stimulation blocks and 60 s rest periods, except for the 1 Hz stimulation, which lasted 20 s. The scan time of these experiments were 5 min 40 s and 6 min 20 s, respectively.

For systematic parameter testing, amplitude effects were first investigated by keeping a constant stimulation frequency of 10 Hz while increasing the amplitude in 50 µA increments from 80 to 230 µA over several experiments. Based on these experiments, the amplitude was then kept constant at 130 µA and the frequency was set to 1, 40 and 100 Hz.

Finally, it was tested whether a prolonged 1 Hz stimulation causes a sustained effect. For this purpose, continuous 1 Hz stimulation at 130 µA for 10 minutes was performed inside the MRI system but without running fMRI. Block-design experiments with 10 Hz at 130 µA, before and after the 1 Hz stimulation were used as readout.

### Transcranial perfusion and immunohistochemistry

At the end of the experiments, mice were deeply anesthetized and transcardially perfused with 4% paraformaldehyde in 0.1 M phosphate buffer. Following dissection, brains were sectioned on a vibratome and processed for immunohistochemistry to verify hippocampal sclerosis and electrode positions.

### Statistical Analysis

Statistical analysis was performed using FSL (v5, FMRIB, Oxford, UK) and Matlab (2016b, Mathworks, USA). All fMRI activations were displayed using cluster thresholding z = 3.1 and p = 0.001 (for implementation we refer to the user guide of the FSL function “FEAT”). Two control mice had to be excluded because the ground/reference electrodes were not connected to the brain. One kainate mouse died during the period of the experiments. All statistical tests and animal numbers are given in the results and figure captions.

## 3. Results

The stimulations were applied in the septal pole of the ipsilateral HC. Based on our previous study [28], hippocampal sclerosis on this side was identified in kainate-injected mice by a slight signal increase in the T2-weighted reference images and an enlarged dentate gyrus due to granule cell dispersion (Fig. 1A).

**Figure 1:**
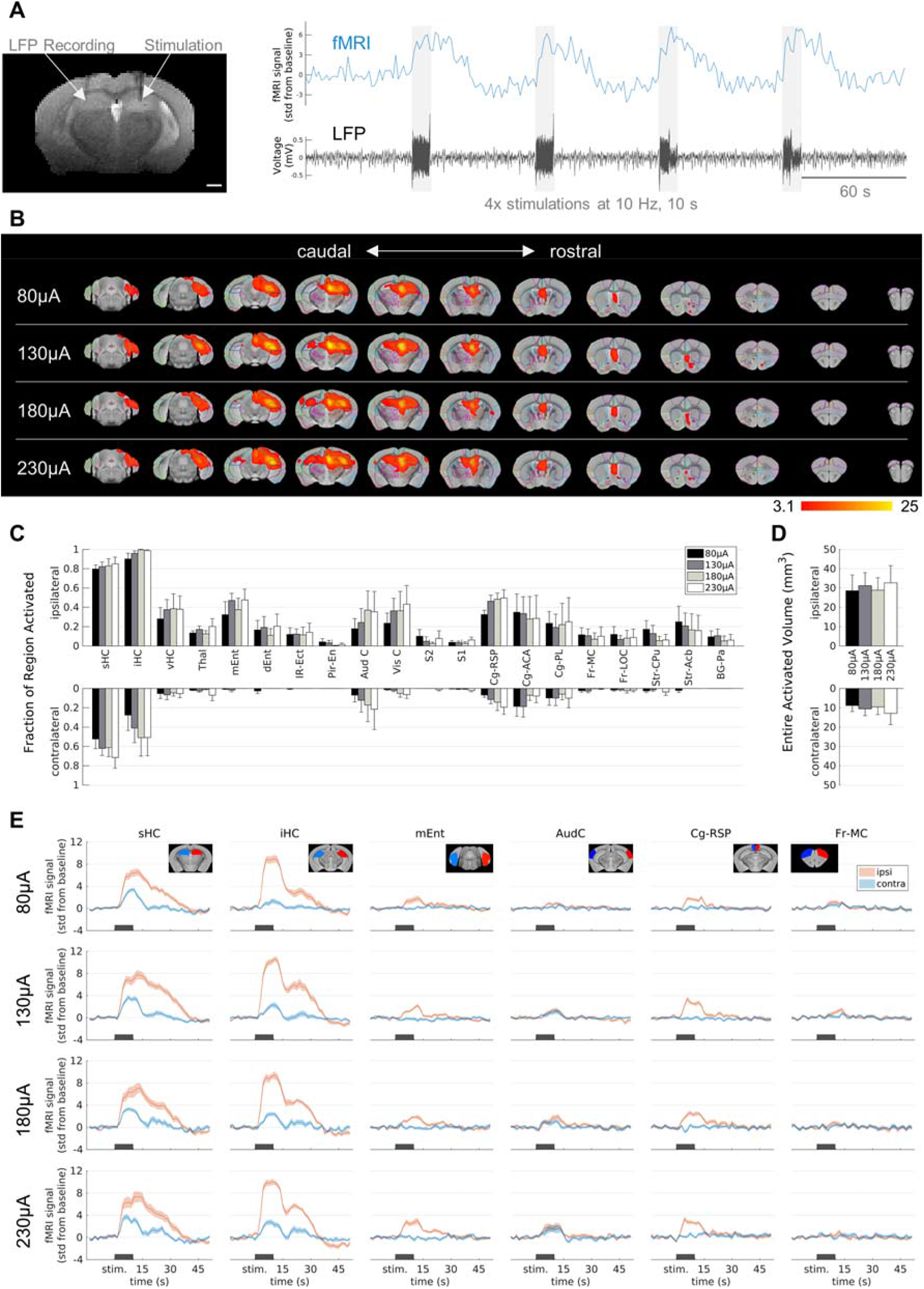
Influence of the stimulation amplitude on fMRI activation in epileptic mice. **A)** Experimental design. Left) Representative T2-weighted anatomical reference image showing stimulation and recording electrodes implanted in the right and left hippocampus, respectively. Scale bar: 1 mm. Right) Exemplary fMRI time course (blue) showing the responses to the stimulation indicated by the electrical artifacts in the LFP trace (black). Scale bar: 60 s. **B)** fMRI activation maps for stimulation amplitudes of 80 µA (top row) to 230 µA (bottom row) from caudal (left) to rostral (right) direction. The group-level fMRI activations are shown as red-yellow overlays in the AMBMC reference space (gray background) with contours of the AMBMC mouse brain atlas* (colors red to yellow represent z-scores from 3.1 to 25 of significant responses at a cluster-defined threshold p <0.001; group n = 6 for 80 and 130 µA, n = 4 for 180 and 230 µA). **C)** Quantification of the activated fractions in atlas regions for stimulation amplitudes from 80 to 230 µA (mean ± standard error of the mean, SEM). Increasing the amplitude did not result in significant differences of the activated volumes (one-way ANOVA p >0.05 with Tukey-Kramer multiple comparison tests). **D)** Quantification of the entire activated volume on the ipsilateral and contralateral hemisphere (mean ± SEM). No significant differences were found between the stimulation amplitudes (one-way ANOVA p >0.05 with Tukey-Kramer multiple comparison tests). However, the activated volumes on the ipsilateral hemisphere were significantly larger than on the contralateral side (one-sample t-tests between ipsi-and contralateral ‘Entire Activated Volumes’; Bonferroni-Holm corrected p-values = 0.019, 0.017, 0.013 and 0.015 for 80, 130, 180 and 230 µA, respectively). **E)** Mean fMRI responses in selected regions (mean ± SEM). The black bars indicate the stimulation periods. *Atlas regions: sHC – septal hippocampus; iHC – intermediate hippocampus; vHC – ventral hippocampus; Thal – thalamus; mEnt – medial entorhinal cortex; dEnt – dorsal entorhinal cortex; IR-Ect – insular region / ectorhinal area; Pir-En – piriform cortex / endopiriform claustrum; Aud C – auditory / temporal region; Vis C – visual cortex; S2 – secondary somatosensory cortex; S1 – primary somatosensory cortex; Cg-Rsp – cingulate region / retrosplenial area; Cg-ACA – cingulate region / anterior cingulate area; Cg-PL – cingulate region / prelimbic area; Fr-MC – frontal association / motor cortex; Fr-LOC – frontal / lateral orbital cortex; Str-CPu – striatum / caudate putamen; Str-Acb – striatum / accumbens; BG-Pa – basal ganglia / pallidum.

### 3.1 Influence of the stimulation amplitude on fMRI activation in epileptic mice

First, the stimulation frequency was kept constant at 10 Hz and the effect of different amplitudes were investigated in chronically epileptic mice. Since preliminary tests at a stimulation amplitude of 30 µA showed only minimal stimulation artifacts in the LFP and no fMRI response (not shown), we increased the amplitude in 50 µA increments. Amplitudes ranging from 80 to 230 µA elicited robust fMRI activations (Fig. 1B).

At 80 µA, the main activation was around the stimulation site in the ipsilateral septal HC (Fig. 1B, first row). It was restricted to the dorsal to intermediate HC and did not expand into the ventral part of the HC. In caudal direction, activation affected parts of the entorhinal cortex and in rostral direction, parts of the cingulate cortex and the septal area. Responses at higher amplitudes (130-230 µA) showed approximately the same activation patterns with only a slightly wider spread (Fig. 1B, second to fourth row).

To quantify the extent of activation, the percentage of the area covered by the fMRI activation was calculated in each brain region (Fig. 1C). Group comparisons across the different brain regions revealed no significant differences between the different amplitudes (one-way ANOVA p >0.05 with Tukey-Kramer multiple comparison test). The entire activated volume did also not change significantly with amplitude (Fig. 1D).

The contralateral septal and intermediate HC became activated only at higher amplitudes (Fig. 1B). However, the fMRI time courses of these areas illustrate that there was already a weak response at 80 µA and only minor changes occurred at higher amplitudes (Fig. 1E).

Overall, the activity patterns were very similar across all current amplitudes. The activated volumes on the ipsilateral hemisphere were significantly larger than on the contralateral side (one-sample t-tests between ipsi- and contralateral ‘Entire Activated Volumes’; Bonferroni-Holm corrected p-values = 0.019, 0.017, 0.013 and 0.015 for 80, 130, 180 and 230 µA, respectively, Fig. 1D).

### 3.2 Influence of the stimulation frequency on fMRI activation in epileptic mice

Next, the amplitude was kept constant at 130 µA and the frequencies 1, 40 and 100 Hz were examined in chronically epileptic mice (Fig. 2).

**Figure 2:**
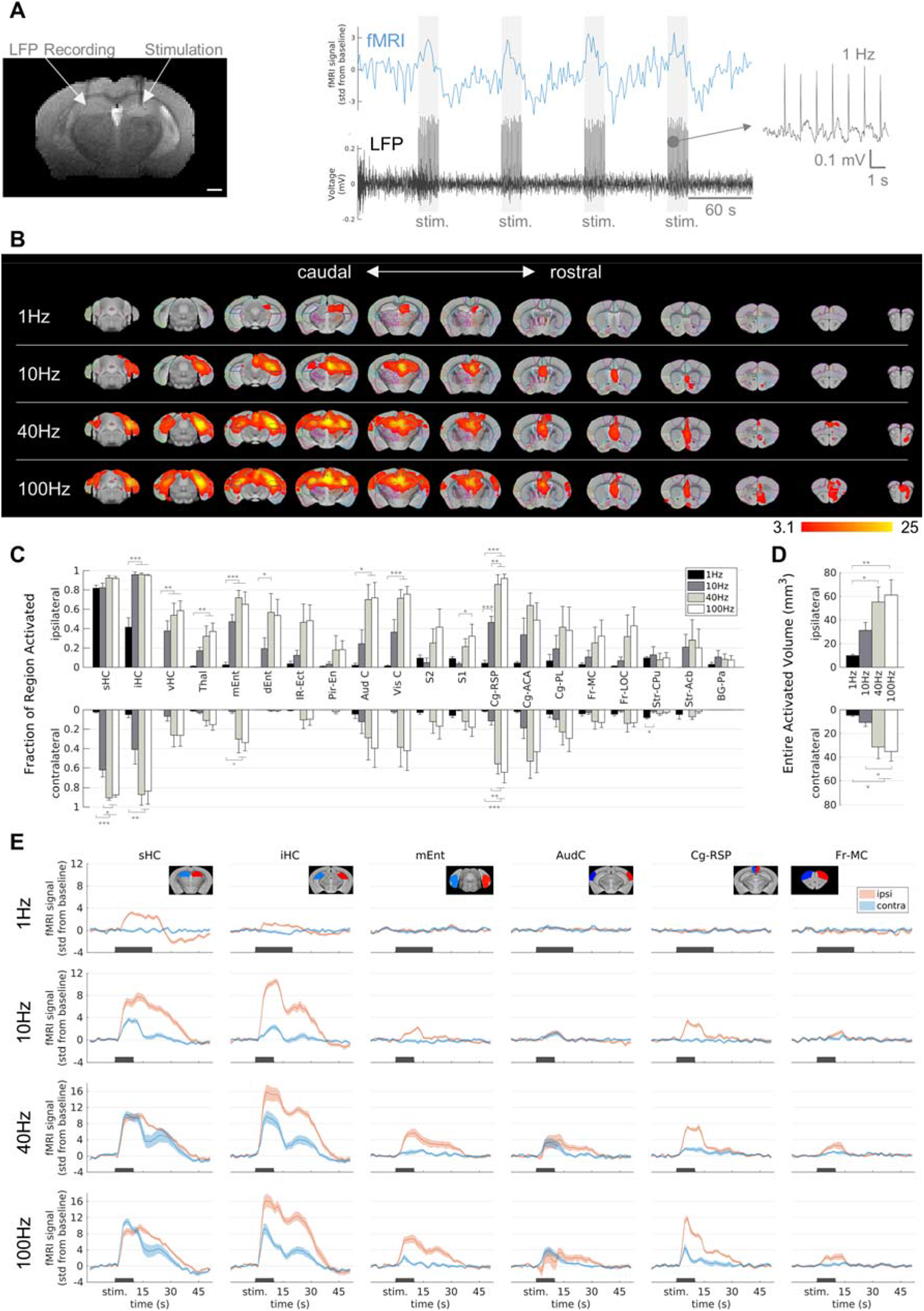
Influence of the stimulation frequency on fMRI activation in epileptic mice. **A)** Experimental design. **B)** fMRI activation maps for stimulation frequencies of 1 Hz (top row) to 100 Hz (bottom row) from caudal (left) to rostral (right) direction. The group-level fMRI activations are shown as red-yellow overlays in the AMBMC reference space (gray background) with contours of the AMBMC mouse brain atlas (colors red to yellow represent z-scores from 3.1 to 25 of significant responses at a cluster-defined threshold p <0.001; group n = 6 for 1 and 10 Hz, n = 4 for 40 and 100 Hz). **C)** Quantification of the activated fractions in atlas regions showed with increasing frequency several significant changes (*p <0.05, **p<0.01, ***p<0.001, one-way ANOVA with Tukey-Kramer multiple comparison tests). **D)** Quantification of the entire activated volume on the ipsilateral and contralateral hemisphere showed significant increases from 1-10 to 40-100 Hz (*p <0.05, **p<0.01, one-way ANOVA with Tukey-Kramer multiple comparison tests). The activated volumes differed significantly between the ipsi-and contralateral hemispheres at frequencies of 1 and 10 Hz, but not at 40 and 100 Hz (one-sample t-tests between ipsi-and contralateral ‘Entire Activated Volumes’; Bonferroni-Holm corrected p-values = 0.017, 0.046, 0.059 and 0.066 for 1, 10, 40 and 100 Hz, respectively. **E)** Mean BOLD fMRI responses in selected regions (mean ± SEM). The black bars indicate the stimulation periods.

At 1 Hz, activation was visible only around the stimulation site, whereas with increasing frequency, activation spread from the HC to large parts of the brain (Fig. 2B). At 100 Hz, fMRI responses were detected in the thalamus, several cortical, cingulate and frontal regions. The activated fractions in several brain regions (Fig. 2C) and the total activated volumes increased significantly with frequency (Fig. 2D, one-way ANOVA p <0.05 with Tukey-Kramer multiple comparison tests).

Another noteworthy change from 10 Hz to 40-100 Hz was the stronger recruitment of the contralateral hemisphere, for example in the intermediate HC, the medial entorhinal cortex, and the cingulate areas. The activated volumes differed significantly between the ipsi-and contralateral hemispheres at frequencies of 1 and 10 Hz, but not anymore at 40 and 100 Hz (one-sample t-tests between ipsi- and contralateral ‘Entire Activated Volumes’; Bonferroni-Holm corrected p-values = 0.017, 0.046, 0.059 and 0.066 for 1, 10, 40 and 100 Hz, respectively. Fig. 2D).

The fMRI time courses show that the response at 1 Hz was comparatively weak but robust (Fig. 2E). Overall, the evaluated fMRI patterns showed only little inter-animal variability (Supplementary figure 1).

### 3.3 Comparison of the stimulated responses in healthy controls and epileptic mice

To investigate whether the spread of activity was related to alterations in the epileptic brain, we performed stimulation experiments on healthy controls for comparison.

As in the epileptic animals, no fMRI response was detected at 10 Hz and 30 µA in the control animals (not shown). However, the first stimulation block at 80 µA triggered afterdischarges (ADs) in control animals (Fig. 3A,B). Subsequently, another block design experiment was performed in which no ADs occurred (Fig. 3C,D). Here, fMRI activation was detected bilaterally in the septal HC, in parts of the entorhinal cortex, cingulate cortex and septum. In contrast to the epileptic mice, activations on the ipsi- and contralateral hemispheres were approximately symmetrical.

**Figure 3:**
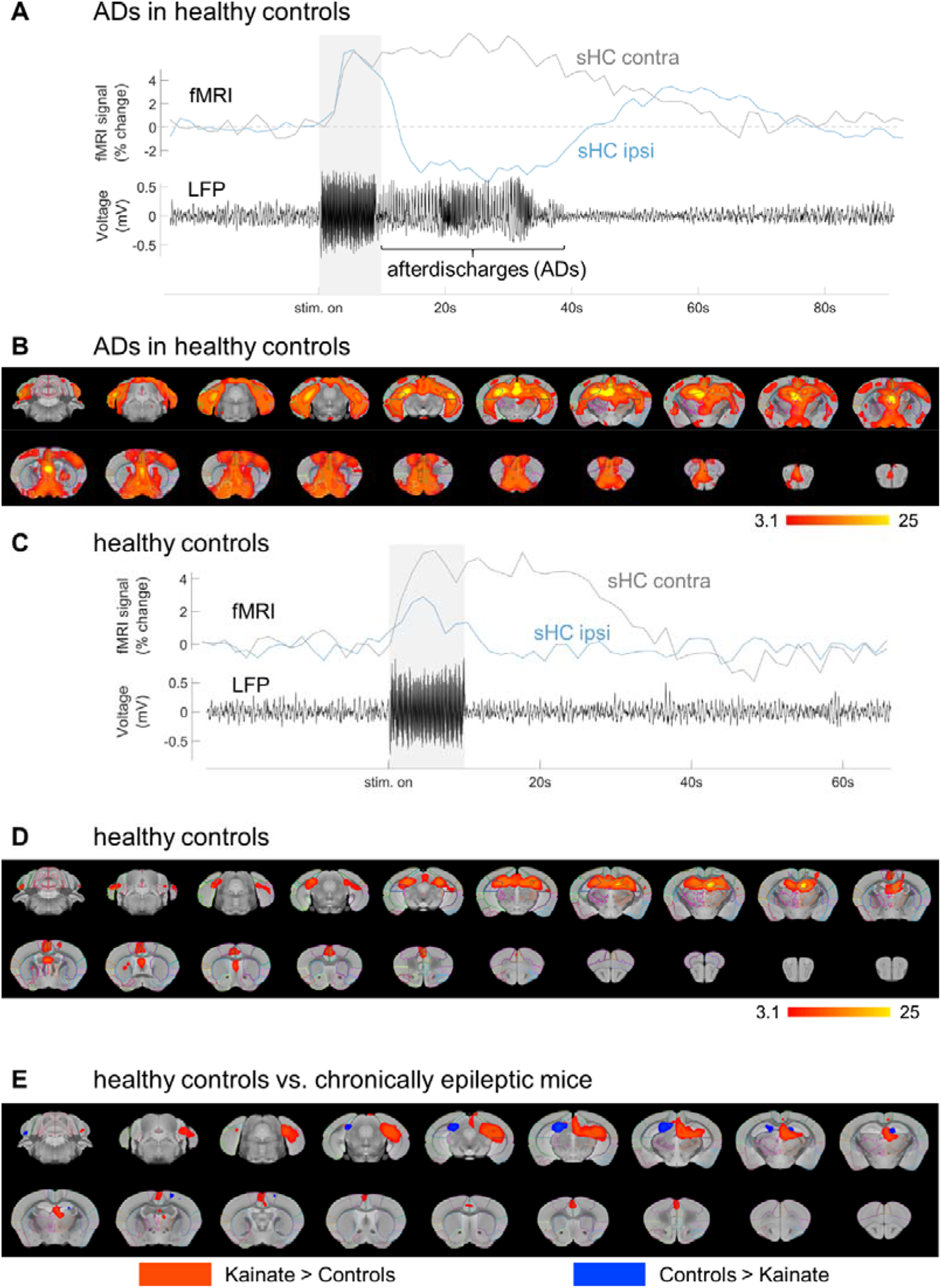
Comparison of the stimulated responses in healthy controls and epileptic mice. **A)** In control mice, LFP recordings in the contralateral HC (black trace) revealed afterdischarges (ADs) triggered by the first stimulation (shaded area). Representative fMRI time courses from locations in the ipsilateral (blue) and contralateral (gray) septal hippocampus. The ADs occurred in all control mice at the first simulation with 10 Hz at 80 µA. **B)** Group-level fMRI activation map of the ADs triggered in control mice (n = 4, cluster-defined threshold p <0.001). **C)** Representative fMRI (blue) and LFP trace (black) of a control mouse from the following experiments after the ADs. No ADs were elicited in these trials. **D)** Group-level fMRI activation map for stimulation with 10 Hz at 80 µA in control mice without ADs (colors red to yellow represent z-scores from 3.1 to 25 of significant responses at a cluster-defined threshold p <0.001; group n = 4). **E)** Group-level differences between epileptic (n=6) and control (n=4) mice (without ADs). Red colors indicate areas in which epileptic animals showed stronger responses, whereas blue colors show areas in which controls showed stronger responses (unpaired two-group difference, cluster-defined threshold p <0.001, stimulations at 10 Hz and 80 µA).

Group differences between healthy and epileptic animals revealed a stronger response in the contralateral septal HC and in a small part of the ipsilateral septal HC for control animals (Fig. 3E). The epileptic animals showed a stronger response in the ipsilateral intermediate HC and in parts of the ipsilateral thalamus, entorhinal cortex and cingulate cortex.

Note that ADs occurred almost exclusively in control animals. Stimulations in epileptic animals resulted in a seizure only in one animal at 40 Hz and 130 µA, a repetition on another day also led to a seizure. In contrast to the control mice, the seizures in the epileptic mouse were associated with strong seizure-induced movements. This precluded further analyses of the fMRI signal and these data sets had to be excluded.

### 3.4 Influence of prolonged 1 Hz stimulation on excitability

We tested if prolonged 1 Hz stimulation affects excitability. We therefore compared the fMRI patterns induced by 10 Hz block-design fMRI before and after 1 Hz stimulation (Fig. 4A).

**Figure 4:**
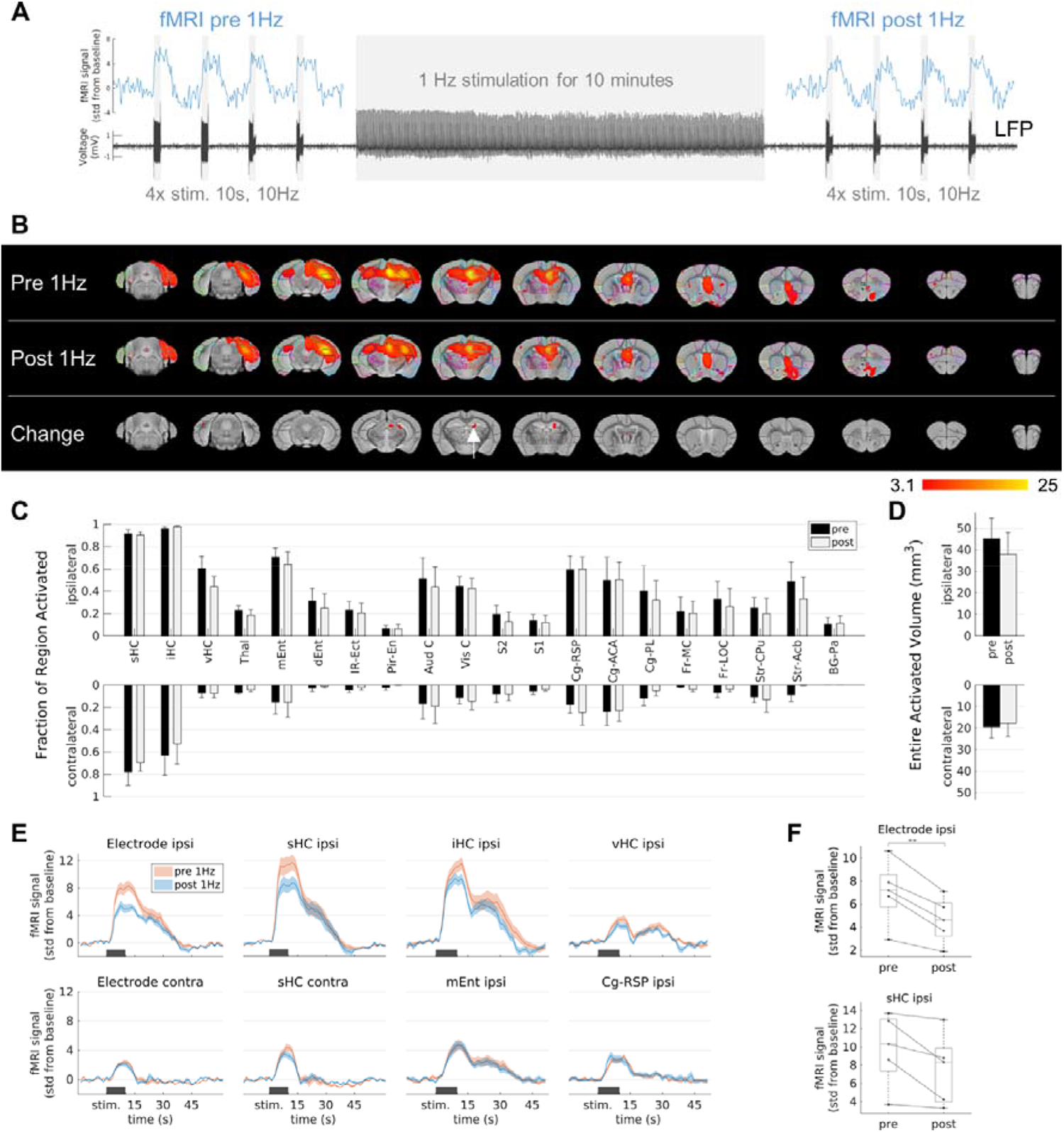
Influence of prolonged 1 Hz stimulation on excitability. **A)** Experimental design. Stimulation at 1 Hz was applied for 10 minutes and block-design fMRI experiments (4 stimulation blocks for 10 s at 10 Hz and 130 µA) were performed before and after the 1 Hz stimulation to evaluate the activation patterns. **B)** fMRI activation maps before (first row, ‘Pre 1 Hz’), after the 1 Hz stimulation (second row, ‘Post 1 Hz’), and the group-level comparison between them (third row). A significant decrease (two-sample paired t-test, cluster-defined threshold p <0.001; group n = 5) was detected in a small area (red color) within the ipsilateral septal HC (white arrow). **C)** Quantification of the activated fractions in atlas regions (mean ± SEM). There were no significant changes from pre 1 Hz to post 1 Hz (paired t-tests p >0.05). **D)** Quantification of the entire activated volumes on the ipsilateral and contralateral hemisphere (mean ± SEM). There was no significant change from pre 1 Hz to post 1 Hz (paired t-tests p >0.05). **E)** Average fMRI responses in selected regions (mean ± SEM). The fMRI response after the 1 Hz stimulation (blue) was reduced compared to the fMRI response before the 1 Hz stimulation (red) in the regions-of-interest (ROIs) near the stimulation electrode (ROIs ‘Electrode ipsi’ and ‘sHC ipsi’), whereas no change was observed in ROIs far from the stimulation site (e.g. at the contralateral side or in the entorhinal cortex and cingulate region). The ROIs ‘Electrode ipsi’ and ‘Electrode contra’ correspond to a volume of 0.5 mm^3^ (eight voxels) directly around the electrodes. These and the ROIs in the septal HC (‘sHC ipsi’ and ‘sHC contra’) were manually drawn in the single subject space to exclude the electrode artefacts. **F)** Change of fMRI amplitudes in the ROIs ‘Electrode ipsi’ (paired t-test between pre and post 1 Hz: **p<0.01) and ‘sHC ipsi’ (paired t-test: p 0.06). Mean fMRI signals in a 10 s window shifted by 5 s from the start of the stimulation to account for the delayed fMRI response.

The extent of the fMRI pattern before 1 Hz stimulation was similar to that after 1 Hz stimulation (Fig. 4B). However, the group comparison revealed a significant decrease within the ipsilateral septal HC (Fig. 4B, third row). There were no significant changes in the activated fraction in individual regions (Fig. 4C) or in the total volumes (Fig. 4D).

To further clarify this finding, the fMRI time courses before and after the 1 Hz stimulation were compared in volumes directly around the electrode, in the entire septal, intermediate and ventral HC (Fig. 4E). We found that the fMRI response in the areas close to the stimulation electrode were reduced after the 1 Hz stimulation and this effect became weaker with increasing volume and distance from the electrode. More distant regions, such as the entorhinal cortex, the cingulate region or the contralateral HC showed no change. The fMRI response was not only reduced on average, but also in each animal (paired t-tests; p = 0.004 in volume ‘Electrode ipsi’, p = 0.06 in volume ‘sHC ipsi’, Fig. 4F).

## 4. Discussion

This study systematically investigated brain-wide fMRI activation patterns following different stimulation protocols in chronically epileptic and healthy control mice. Importantly, our results were obtained in the ihKA mouse model of mTLE that replicates several key features of the human disease, such as hippocampal sclerosis and recurrent seizures [29] .

We found that the stimulation amplitude has little influence on the extent of the induced fMRI patterns over a relatively wide amplitude range. 80 µA was the lowest amplitude in our study that evoked an fMRI response and higher currents led only to a minor, yet unspecific broadening of the activation pattern, but did not recruit fundamentally new areas. This is in line with a previous clinical study where two current levels (4 and 8 mA) were compared and no difference was observed between the activation patterns [21]. The absolute values of the stimulation current may differ between studies due to differences in electrode design and/or positions, but it can be concluded that if a response to a stimulation is detectable, modifying the current strength is unlikely to result in a significantly different response or functional outcome.

The results of the present study show that the stimulation frequency constitutes the key factor in determining which areas/networks of the brain are affected by a stimulation. The extent of activation could be adjusted from local activation at 1 Hz to brain-wide activation at 100 Hz.

Other studies in healthy rodents have reported that activity induced at 10 Hz in the dorsal HC remained very confined to the dorsal part of the HC and concluded that frequencies above 20 Hz do not result in a polysynaptic spread [22,23,53]. Only when stimulating the intermediate HC or after induced long-term potentiation (LTP), a more widespread activation was detected [54–56]. Our results from the ihKA mouse model of TLE showed upon 10 Hz stimulation, a spread to the intermediate HC, entorhinal cortex, cingulate and septal regions. The response in these regions was also stronger compared to the healthy controls, indicating increased functional connectivity in the epileptic brain, which has also been found in rodent models of TLE by resting-state fMRI [57,58]. Stimulations at 40 and 100 Hz elicited clear responses in cortical, frontal and prefrontal areas. This is particularly important because the propagation of activity to frontal areas, which are important in the control of executive functions, may constitute motor dysfunction during epileptic seizures [59].

One surprising result of this study was that the stimulations triggered epileptiform ADs in healthy controls, whereas in epileptic animals, seizures occurred in only one animal. This represented only 4.1 % (2 of 49) of all stimulation experiments in epileptic mice, and was therefore not representative. It is still unknown what exactly triggers epileptic seizures. The seizures themselves, in epileptic animals as well as patients, are difficult to study with fMRI because they occur spontaneously and are often associated with strong movements. The fact that the stimulations elicited epileptiform ADs only in healthy controls and that these animals did not move during the ADs, suggests that the mechanisms of ADs in healthy controls and spontaneous epileptic seizures are different. The results from studies of stimulated ADs in healthy animals, e.g. [42,44,45,60], therefore cannot necessarily be generalized to real epilepsy.

Aiming to interfere with epileptiform activity, recent non-imaging studies explored the effect of 1 Hz stimulation in mTLE patients [61] and the ihKA mouse model of mTLE [36]. The underlying strategy is that prolonged stimulation leads to reduced excitability, which may be associated with reduced seizure incidence. We measured a reduced fMRI response after prolonged 1 Hz stimulation supporting this strategy to reduce excitability. The effect was relatively weak, but the applied stimulation duration of only 10 minutes can be extended to further reduce the excitability.

A limitation of our work is that stimulations were applied only during fMRI measurements. However, a stimulation aimed at reducing epileptic activity would require long-term application, which may lead to alterations over time. A future study should investigate the effects of long-term stimulation on epileptic activity and examine changes in induced activity at multiple time points using fMRI.

Another limitation is that this study was performed only in an animal model of mTLE. In the human disease, there is greater variability between patients, the epileptic foci might not be so restricted and there is no guarantee that the electrode is positioned exactly at the focus. This variability could lead to suboptimal stimulation parameters. For example, our results showed that 1 Hz stimulation elicits only a local response, which may be beneficial to minimize side effects [62]. However, to effectively interfere with the occurrence of epileptic activity, a therapeutic stimulation must reach the epileptic focus. If this is not achieved by the local effect of 1 Hz stimulation, it may be necessary to increase the stimulation frequency. Controlling stimulation responses using fMRI may therefore provide a way to more effective, individualized treatment.

## 5. Conclusions

We here present the brain-wide responses to hippocampal stimulation in the ihKA model of mTLE. The key parameter that determined the extent of activation was the stimulation frequency. While low frequency (1 Hz) stimulation elicited only a local response, high frequency (40-100 Hz) stimulation resulted in spread of activity from the HC to cortical and frontal regions in the epileptic brain. Compared to healthy controls, epileptic mice showed stronger activity in the intermediate HC, entorhinal and septal region, indicating increased functional connectivity to the stimulation site in the sclerotic HC.

The amplitude parameter, in contrast to frequency, offers little opportunity to vary the stimulation effects. However, the fact that the amplitude did not show a large influence makes it possible to compare the results of studies even if the amplitude settings differ. This supports the translational value of preclinical fMRI studies. It remains to be investigated whether fMRI also shows plastic changes and altered activity patterns in the case of a chronic application of stimulation.

## CrediT authorship contribution statement

**Niels Schwaderlapp:** Conceptualization, Methodology, Software, Formal analysis, Investigation, Data Curation, Writing – Original Draft, Writing – Review & Editing, Visualization. **Enya Paschen:** Conceptualization, Methodology, Investigation, Data Curation, Writing – Review & Editing. **Pierre LeVan:** Writing – Review & Editing, Funding acquisition. **Dominik von Elverfeldt:** Resources, Writing – Review & Editing, Project administration, Funding acquisition. **Carola Haas:** Resources, Writing – Review & Editing, Supervision, Project administration, Funding acquisition.

## Abbreviations

Ads: afterdischarges
DBS: deep brain stimulation
HC: hippocampus
ihKA: intrahippocampal kainate
LFP: local field potential
mTLE: mesial temporal lobe epilepsy

