## Supplementary Figure 1 for "Probing hippocampal stimulation in experimental temporal lobe epilepsy with functional MRI"

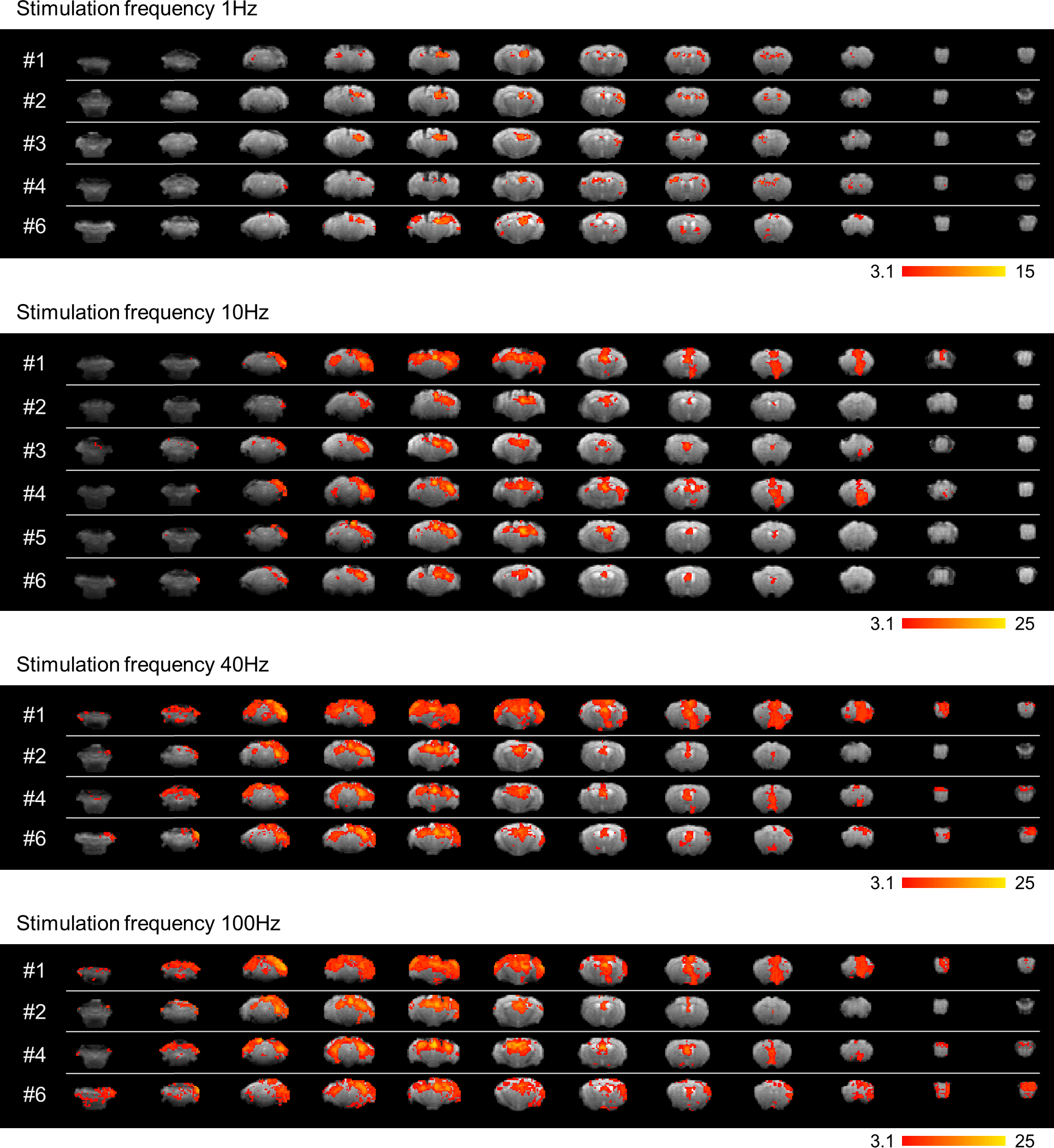

**Supplementary figure 1: Individual fMRI activation maps of kainate mice**

fMRI activations (at 130 µA, 1-100 Hz) of kainate mice in their original field-of-view (without registration to the AMBMC reference space). The fMRI activation was overlaid (colors red to yellow) on an fMRI volume of this experiment. One mouse (#5) died during the period of the experiments (but not during or closely after an fMRI measurement), and one mouse (#3) exhibited epileptic seizures at 40 Hz and was therefore not included in the group analysis of 40 and 100 Hz.
